# Kinship does not predict the structure of a shark social network

**DOI:** 10.1101/853689

**Authors:** Johann Mourier, Serge Planes

## Abstract

Genetic relatedness in animal societies is often a factor that drives the structure of social groups. In the marine world, most studies which have investigated this question have focused on marine mammals such as whales and dolphins. For sharks, recent studies have demonstrated preferential associations among individuals from which social communities emerge. Assortment patterns have been found according to phenotypic or behavioural traits but the role of genetic relatedness in shaping the social structure of adult shark populations has, to the best our knowledge, never been investigated. Here, we used a social network analysis crossed with DNA microsatellite genotyping to investigate the role of the genetic relatedness in the social structure of a blacktip reef shark (*Carcharhinus melanopterus*) population. Based on data from 156 groups of sharks, we used generalized affiliation indices to isolate social preferences from non-social associations, controlling for the contribution of sex, size, gregariousness, spatial and temporal overlap on social associations, to test for the influence of genetic relatedness on social structure. A double permutation procedure was employed to confirm our results and account for issues arising from potentially elevated type I and type II error rates. Kinship was not a predictor of associations and affiliations among sharks at the dyad or community levels as individuals tended to associate independently of the genetic relatedness among them. The lack of parental care in this species may contribute to the breakdown of family links in the population early in life, thereby preventing the formation of kin-based social networks.

## INTRODUCTION

Group formation is an adaptive strategy, widespread across the animal kingdom, that can take various forms, from temporary unstable associations to long-term stable groups in complex societies (Krause and Ruxton 2002). Understanding the factors that influence the formation and evolution of social groups is important in order to understand the evolution of animal societies as well as to gain insight into population dynamics and to inform conservation strategy (Snijders et al. 2017). Associations among individuals can provide benefits to improve individual fitness by, for example, reducing predation risk or improving foraging efficiency (Krause and Ruxton 2002). While individuals can benefit by simply associating with other conspecifics (e.g., Kerth et al. 2011), the benefit of grouping can be enhanced by associating with similar individuals, also called social assortativity. By associating with individuals of the same size or the same sex, individuals are more likely to avoid conflict or harassment (Dadda et al. 2005) and their risk of predation is reduced via the confusion effect (Landeau and Terborgh 1986). Further, assorting with kin can also provide indirect fitness benefits (Hamilton 1964). Kin assortment has been shown to provide benefits in reducing aggression (Olsén and JäUrvi 2005) or increasing growth rate (Brown and Brown 1993).

Kin structuring has received extensive attention in many animal societies, especially where animals form stable breeding groups or where groups arise from the retention of offspring and delayed dispersal that facilitates the development of interactions with relative and kin-based groups (Wolf and Trillmich 2008; Hatchwell Ben J. 2010; Wiszniewski et al. 2010). In groups composed of relatives, kin selection should play a role in determining cooperation among group members (Hamilton 1964), although cooperation can arise also between non-kin or when only a subset of the group is related (Clutton-Brock 2009). The role of relatedness in structuring animal societies that are characterised by a dynamic fission-fusion social system has been well studied in species with parental care such as dolphins, giraffes, elephants or bats (Wittemyer et al. 2009; Wiszniewski et al. 2010; Kerth et al. 2011; Carter et al. 2013), but much less is known for species without parental care, as is the case for many species of fish (<u>but see</u> Croft et al. 2012). While the link between social networks and kinship has been extensively studied in terrestrial animals (Holekamp et al. 2012; Carter et al. 2013; Arnberg et al. 2015), kinship structure in social networks of marine and freshwater organisms has been primarily limited to marine mammals (Wiszniewski et al. 2010; Mann et al. 2012; Reisinger et al. 2017). Several cetacean societies show strong kin-biased social network structures. However, in fishes, kin structure is less clear. Work on guppies, for example, did not find kin assortment, even in species that are capable of kin discrimination (Croft et al. 2012). While sharks have recently been shown to be able to develop preferred associations and organise into structured social networks (Mourier et al. 2018), kinship has only been explored in one case study that focused on juvenile sharks (Guttridge et al. 2011) but did not find any clear influence of kinship in association patterns even for juvenile sharks, highlighting a lack of information on the potential for kin-based associations to arise in shark populations. Another study on spotted eagle rays did not find any evidence of relatedness in the formation of groups (Newby et al. 2014), although association strength was not quantified using association indices.

Overall, most studies that have explored the relationship between genetic relatedness and social interactions have focused on highly social species and in particular, on species that exhibit parental care (Wolf and Trillmich 2008; Wiszniewski et al. 2010; Kerth et al. 2011). Studying less social vertebrates should significantly improve our understanding of how social and genetic structure interact to shape the evolution of sociality in the animal kingdom.

In this study, we investigate the interaction between socio-spatial patterns and genetic relatedness in a population of blacktip reef sharks (*Carcharhinus melanopterus*) monitored over a 3-year period on the north shore of Moorea Island (French Polynesia). Sharks represent an interesting and unique model to explore the extent to which individuals interact with genetically related associates due to ecological traits that differ from most social vertebrates. Like most social animals, sharks are now increasingly recognised as being capable of complex social interactions, developing preferred social associations (Guttridge et al. 2009; Jacoby et al. 2010; Mourier et al. 2012), showing unexpected learning abilities (Guttridge et al. 2013; Mourier et al. 2017) and developing patterns of leadership and dominance hierarchy (Guttridge et al. 2011; Jacoby et al. 2016; Brena et al. 2018). However, contrary to many social organisms, reef sharks do not show parental care and almost all shark species drop their progeny in specific nurseries outside adult habitats and leave them to interact by themselves (Mourier and Planes 2013). These discrete nurseries are chosen to potentially provide the neonates with a safe environment where they will spend their first months of life. Recent studies suggested that females show reproductive and even natal philopatry to these particular birthing grounds (Mourier and Planes 2013; Feldheim et al. 2014), suggesting that newborn sharks may have the opportunity to develop strong relationships with close kin. When juvenile sharks reach a certain size or age, they leave their nursery to explore a wider home range (Chapman et al. 2009); they then integrate within the adult population and start interacting with older individuals, but it is not known whether they coexisted with other newborn during their juvenile stage or disperse alone. Therefore, while aggregations of kin are possible during the early stages, it is currently unknown if they persist through adulthood after dispersal. In shark populations, interactions between kin are also diluted by the presence of numerous neighbours and average relatedness quickly drops with increasing group size (Lukas et al. 2005). In some shark species, the likelihood of associating with a related peer is reduced due to small litter size and a high mortality rate at the juvenile stage, leading to a lack of first order relatives to reach adulthood. However, in a closed system, such as an isolated island, and in the case of blacktip reef sharks which spend their entire life cycle within Moorea (Mourier and Planes 2013), relatives will have more chances to encounter each other and to interact in social groups. Thus, in these conditions, the limited number of related pairs might decrease the risk of inbreeding.

To understand the assortative forces which underpin the structural properties of the system is challenging for elusive underwater animals. As the blacktip reef shark displays a high degree of site fidelity (Papastamatiou et al. 2009) and shares some of its areas with many conspecifics (Mourier et al. 2012), exploring this network holds the potential to work out the relationship between spatial, social and genetic structure in a reef shark. Size, sex and gregariousness of sharks have been shown to influence assortment at the population and community levels (Mourier et al. 2012; Mourier et al. 2017). However, whether genetic relatedness plays a role in structuring the network at both the individual and community levels remains unknown. In particular, whether sharks benefit from associating with kin remains unknown as cooperation has not been proven and social foraging may not require associations with kin to improve predation success (Labourgade et al. 2020).

We aim to test whether the social structure of sharks at different scales can be explained by the genetic relatedness between individuals after controlling for non-social structural factors, including space use, temporal overlap, phenotype, and individual gregariousness.

## MATERIAL AND METHODS

### Field observations and data collection

Between 2008 and 2010, observation surveys were conducted along approximately 10km of coastline of the Northern reef of Moorea Island (French Polynesia) (Figure 1). The surveys consisted of 40 min dives (~30 min dedicated to survey) at 7 sites along a 10 km portion of reef (total = 180 dives, site 1 = 20, site 2 = 50, site 3 = 8, site 4 = 33, site 5 = 30, site 6 = 34 and site 7 = 7). Individual blacktip reef sharks were identified by photo-identification, using unique, lifelong colour-shape of the dorsal fin (Mourier et al. 2012).

**Figure 1.**
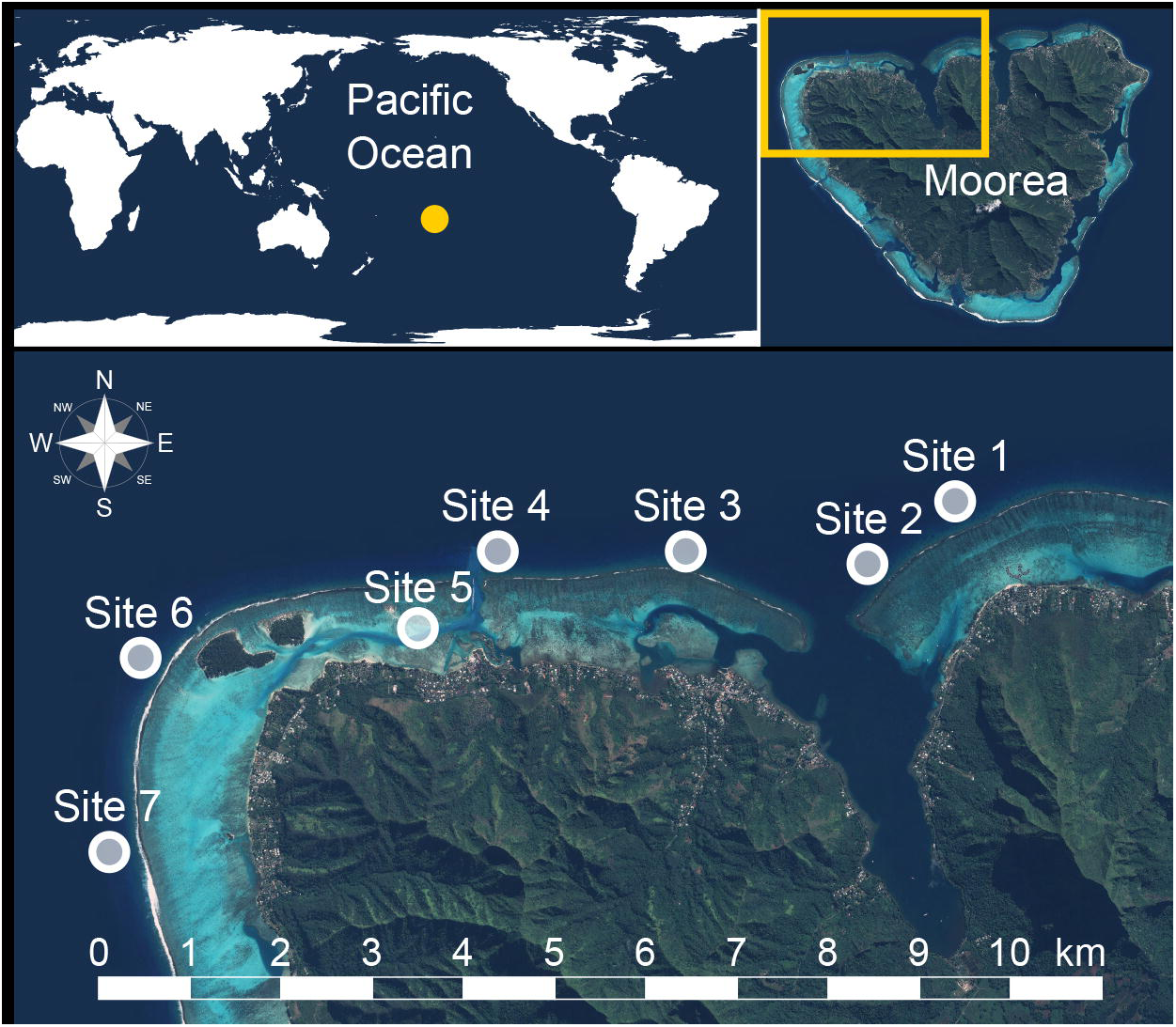
Map of the study location indicating the monitored sites along the 10 km reef edge of the north coast of Moorea.

Associations between individuals were defined using the ”Gambit of the Group” (Whitehead and Dufault 1999) assuming that all individuals observed together are then considered as “associated”. This approach is appropriate when individuals move between groups and direct interactions are difficult to observe, but where groups can be easily defined (Franks et al. 2010; Farine and Whitehead 2015). An experienced diver conducted a stationary visual census at each site monitored, moving and identifying sharks within a ~100 m radius area (made possible by the high visibility conditions in these tropical waters). As most sharks usually remained together during the time of the dive, we considered the largest number of individuals observed within a 10 min period to be part of a group. We are confident that observed associations represented true grouping structure, because groups were spatio-temporally well-defined and sharks were engaged in specific social behaviour (<u>e.g. following, parallel swimming or milling</u>; Mourier et al. 2012). To avoid the potential for weak and non-relevant associations between pairs of individuals with very low number of sightings, we used a restrictive observation threshold to include only individuals observed more than the median number of sightings (median = 14; mean SD = 14.92 8.04; Supplementary Figure S1). Thus, all individuals seen less than 15 times were removed from the analyses to ensure that associations were estimated with high accuracy and precision.

### DNA sampling and laboratory procedures

Shark fishing sessions using rod and reel and barbless hooks were conducted to obtain tissue samples for genetic analysis. Once hooked, sharks were brought alongside the boat where they were inverted and placed in tonic immobility while biological data and tissue samples were collected. Each shark was identified by photo-identification of the dorsal fin, sexed and measured to the nearest centimeter (Mourier et al. 2012; Mourier, Mills, et al. 2013). Fishing sessions were conducted directly after underwater surveys to avoid perturbations of the experimental setup (Mourier et al. 2017) and to increase the chance of getting DNA samples from sharks that were part of the social network. Fishing effort was maintained until sharks failed to respond to the bait (generally <30 min and after catching 2-3 individuals). A fin clip was collected from the second dorsal fin or anal fin and samples were individually preserved in 95% ethanol and returned to the laboratory for genotyping (Mourier and Planes 2013). DNA was extracted using the QIAGEN DX Universal Tissue Sample DNA Extraction protocol. PCR amplification and the microsatellite loci used are described in detail in previous studies (Mourier and Planes 2013; Vignaud et al. 2013; Vignaud et al. 2014). The software MICROCHECKER (Van Oosterhout Cock et al. 2004) was used to test for null alleles and other genotyping errors.

We compared the suitability of seven pairwise relatedness estimators: five non-likelihood estimators (Queller and Goodnight 1989; Li et al. 1993; Ritland 1996; Lynch and Ritland 1999; Wang 2002) and two maximum-likelihood estimators (Milligan 2003; Wang 2007) in the R package *related* (Pew et al. 2015) and determined that the triadic maximum-likelihood estimator (<u>TrioML</u>; Wang 2007) was best suited to our microsatellite panel (Supplementary materials, Supplementary Figure S2) as it showed the highest correlation (i.e. 0.831) with the true values and the smallest variation around the mean for every relationship (except for full-sibs). This analysis generates simulated individuals of known relatedness based on the observed allele frequencies and calculates the genetic relatedness using the different estimators. The correlation between observed and expected genetic relatedness was obtained for each estimator, and the one with the highest correlation coefficient was selected for further analysis.

### Defining associations

Using R package *asnipe* (Farine 2013), we calculated dyadic association strengths (i.e. associations among pairs of individuals) among photo-identified individual sharks seen in groups from the spatio-temporal co-occurrences, as the proportion of time two individuals were observed together at the same site given that at least one was observed, using the simple-ratio association index (SRI) (Cairns and Schwager 1987). The SRI is the recommended association index when calibration data are unavailable (Hoppitt and Farine 2018).

To measure the diversity of associations, we calculated the social differentiation (S) in the network that is the estimated coefficient of variation (standard deviation divided by mean) of the true association indices. If the social differentiation of the network is 0, then relationships among members are completely homogeneous. Conversely, if the social differentiation is above 1.0, there is considerable diversity in the relationships between pairs of individuals within the network (Whitehead 2008). For our data, the standard error of S was generated by bootstrapping (1 000 replications).

### Potential structural factors of social associations

We quantified five structural factors that could affect shark association patterns: spatial overlap, temporal overlap, gregariousness, and size and sex similarity for each pair of individuals. Genetic relatedness was not included as a structural factor as it was tested independently when other factors are extracted.

For each individual, an encounter rate (i.e., no. sightings of individual at site, divided by no. sampling occasions at site) was calculated by site to define individual spatial utilization (Supplementary Figure S3). We then generated a Bray-Curtis similarity matrix of space use to construct a matrix of spatial overlap between individuals using R package “vegan” (Dixon 2003).

Individuals using an area at the same time are more likely to be associated with each other. The study period corresponds to a total of 28 months between February 2008 and June 2010. The temporal overlap was calculated as the SRI calculated on whether pairs were observed in the study area within sampling periods of 60 days.

Gregariousness was calculated following Whitehead and James’s (2015) correction, where the gregariousness predictor between two individuals (*a* and *b*) is the log of the sum of the association indices involving *a* (except the *ab* index) multiplied by the sum of those involving *b* (except the *ba* index):

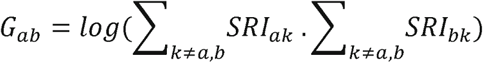

where *SRI*_*ab*_ is the association index between individuals *a* and *b*, and *SRI*_*kk*_ is set to zero for all *k*.

Shark length was classified into size classes ranging from 1 to 6 (1: TL < 110 cm; 2: 110-119 cm; 3: 120-129 cm; 4: 130-139 cm; 5: 140-150 cm; 6: TL > 150 cm). For sex and size similarity, we constructed a binary matrix in which elements a_*ij*_ = 1 when individuals *i* and *j* were of the same class and a_*ij*_ = 0 otherwise (sex class, 1 if same sex, 0 if not; size class, 1 if same size class, 0 if not).

### Influence of structural factors on social associations

We quantified the contribution of all five structural factors in driving social patterns with a multiple regression quadratic assignment procedure (MRQAP) modified by Farine (2013) that enables null models built from pre‐network data permutations to be used in conjunction with a MRQAP regression. This approach was shown to be more accurate than classic MRQAP procedures (Farine 2017). We assessed possible linear relationships between the social associations and the structural factors using the SRI association matrix as the dependent variables and the matrices representing pairwise similarity of each of the five structural factors as independent variables. We used 20,000 permutations to build randomized distributions to compare with the empirical coefficient. The *P-*values were the proportion of the estimated coefficient regression which were smaller or greater than what would have been expected by chance. We used the *mrqap.custom.null* function from *asnipe* R package (Farine 2013) to run MRQAP tests in R v. 3.3.0 (R Core Team 2019).

### Removing the effects of structural factors from associations

We developed generalized affiliation indices (<u>GAI</u>, Whitehead and James 2015) to remove the effects of the significant structural factors from the associations and test the existence of true affiliations between dyads (i.e. active association preferences). For this, we fitted a binomial generalized linear model (GLM) with the unfolded SRI matrix as the dependent variable, and the significant structural factors selected from the MRQAP as independent variables. GAI represents the assortment of individuals not explained by the significant structural factors and corresponds to the deviance residuals of the model. The model was: SRI ~ TO + SO, where SRI is the association matrix, TO is the temporal overlap matrix and SO is the spatial overlap matrix (as only TO and SO were significant factors in the MRQAP, Table 1).

**Table 1:** Multiple Regression Quadratic Assignment Procedure and the influence of all structural variables on shark social associations. Matrices representing structural variables (predictors) were regressed against the association matrix (SRI) using a subset of the individuals in the population (*n* = 43) to which genetic relatedness was available. TO: temporal overlap; SO: spatial overlap; Gregariousness based on Godde et al. (2013); Size and Sex: binary matrices where individuals of the same size/sex classes are represented by 1, and different classes by 0. Adjusted R^2^ indicates how much of the variation on association indices was explained by the predictors. Significant predictors in which *P*-values are given by the proportion of times the empirical regression coefficient was smaller than the null expectancy from 20,000 randomisations.

| Predictors | Regression Coefficient ($\beta$) | $P$ ($\beta \leq r$) | Adjusted $R^2$ |
| --- | --- | --- | --- |
| TO | 0.553 | <0.001* | 94% |
| SO | 0.107 | <0.001* |  |
| Gregariousness | -0.002 | 0.999 |  |
| Size | 0.006 | 0.089 |  |
| Sex | -0.002 | 0.999 |  |

### Social preferences and null models

We used a null model to test both for social preferences and the significance of the observed network modularity. We generated 20,000 randomized association and affiliation networks based on 25,000 data-stream permutations of the raw observation data with a swapping algorithm (Bejder et al. 1998). We permuted the empirical group-by-individual matrix constraining the number of groups, individuals and occurrences (matrix dimension and fill), group size (row totals) and individual frequency of observation (column totals). To minimize the effect of initial values potentially correlated to the empirical data, we removed the first 5,000 randomized matrices. From the randomised group-by-individual matrix, we calculated a simple-ratio index association matrix, with which we built a generalized affiliation index using the same predictors selected via MRQAP for the empirical data. We used a modified version of R codes available from Machado et al. (2019) to build null models and to calculate SRI, GAI and modularity.

We compared the standard deviation (SD) of the observed simple-ratio index (SRI) and the SD of the observed generalized affiliation index (GAI) with the distribution of the SD of corresponding randomized SRI and GAI matrices generated by the null models detailed above. An observed SD significantly higher than the null expectation indicates the presence of a non-random structure in associations and the presence of preferred and avoided associations for affiliations. We also tested for strongly connected social communities by comparing the empirical modularity (Q) (Newman 2006) of SRI and positive GAI matrices with that of the randomized matrices. Social modules (or communities) were determined based on the leading eigenvector of the community matrix (Newman 2006). Empirical SD and Q values were considered statistically significant if they fell above the 95% confidence interval of their randomized distributions.

### Genetic relatedness, social structure and sex differences

To assess whether relatedness differs for same-sex dyads, we constructed three binary matrices (0,1), each encoding the presence of a certain dyad type (female–female, male–male or female–male). We then tested for a correlation with the relatedness matrix using three Mantel tests (20,000 permutations), via the vegan R package (Dixon 2003).

For each type of dyad (female–female, male–male or female–male), we then tested for a correlation between the SRI and GAI matrices and the pairwise genetic relatedness among sharks using Mantel tests and compared the test statistics to those of the 20,000 permuted networks.

We also compared the gregariousness of individual sharks between the sexes. For this, we used two measures of gregariousness: node degree (or binary degree) which is the number of direct neighbours each individual is connected to in the network and node strength (or weighted degree) that is the sum of associations of an individual. These metrics were calculated from the SRI network. We then used these network metrics in order to determine whether males and females differed in their gregariousness. We constructed generalized linear models (GLMs) to test how sex affected the observed network degree (degree ~ sex) and strength (strength ~ sex). We ran these same models with randomized permutations of the network data to evaluate statistical significance (Farine and Whitehead 2015; Farine 2017).

To determine whether individuals within size class and social modules (i.e. network communities) were more or less closely related than expected, we compared the observed values for each size class and social module against a distribution of expected relatedness values generated by randomly shuffling individuals between size classes and social modules for 1000 permutations, where size was kept constant, using the R package *related* (Pew et al. 2015). If the observed mean relatedness was greater than that of the permuted data, then the null hypothesis which predicted that the mean within‐module or within size class relatedness is random, was rejected.

If only a few closely related individuals were present, then it is possible that their within-community overabundance compared to between social communities might not be detected using mean coefficient of relatedness (Buston et al. 2009). In turn, we verified whether the proportion of closely related pairs was higher within than between social communities using a chi-squared test following the same approach as the preceding analysis with mean relatedness. We compared the χ^2^ statistics of the observed difference in proportions of relatedness values above a certain threshold between within- and between-communities to that of expected relatedness values generated by randomly shuffled individuals between community groups for 1000 permutations and keeping size constant. We tested with a threshold relatedness value of 0.25 corresponding to the theoretical relatedness of half-sibs.

Recent studies found that (datastream and node) permutation tests, which are widely used to test hypotheses in animal social network analyses, can produce high rates of type I error (false positives) and type II error (false negatives) (Franks et al. 2020; Puga□Gonzalez et al. 2020; Weiss et al. 2020; Farine and Carter 2020). In order to ensure that the results provided by our GAI analyses are reliable and account for both social and non-social nuisance effects, we used an approach proposed by Farine and Carter (2020) that uses datastream permutations to control for nuisance effects, then uses node permutations to test for the statistical significance of the effect of interest. It first uses datastream permutations to calculate the deviation of each of a node-level or edge-level metric from its random expectation given the structure of the observation data (i.e. equivalent of a residual value), which are then fitted into a model such as an MRQAP or a regression, to generate a corrected test statistic with node permutations used to calculate the significance of this statistic. This approach is similar to generalised affiliation indices (Whitehead and James 2015), but it uses datastream permutation tests, rather than regression models, to estimate the deviance from random. This approach demonstrated its robustness to test the role of kinship in shaping the strength of interactions between individuals in the presence of other social effects such as the presence of non-kin social bonds (Farine and Carter 2020). We therefore applied this double-permutation procedure to test for the role of kinship in driving associations among sharks as well as the role of sex on node centrality (controlling for the number of observations) and to confirm the robustness of our previous results to high type I and type II error rates.

## RESULTS

### Data summary

Of 241 catalogued sharks (150 males, 91 females; Mourier et al. 2012), 49 (36 males, 13 females) were observed on 15 or more occasions (mean resightings = 14.92 ± 8.04 SD, Supplementary Figure S1). A total of 225 adult sharks were genotyped from the studied area. From the 49 sharks included in our social network analysis, 87% (43) were genotyped. Therefore, 43 individuals (30 males, 13 females) were included in the remaining analyses. This resulted in 156 observed groups (mean group size = 8.60 ± 4.92 SD). From the 17 microsatellite markers taken from our previous study (Mourier and Planes 2013), the presence of null alleles was detected at Cli12 which was then removed from our dataset for further genetic analyses. We conducted the genetic analyses with 16 loci (Supplementary Table S1).

### Social structure

The social differentiation of the population was higher than 1 (S ± SE = 1.474 ± 0.037), revealing a diverse range of associations and a well-differentiated society. The most significant predictors of shark associations were the temporal and spatial overlaps, which explained 94% of the total variance in SRI (MRQAP results, Table 1).

We rejected the null hypothesis that sharks associate randomly, as the observed SD of SRI was higher than the distribution of random SD values. When GAI removed the influence of temporal and spatial overlaps from SRI, we also rejected the null hypothesis of random affiliations (Figure 2a) demonstrating the presence of preferred social affiliations. At the population level, the modularity (Q) of the association (SRI) and affiliation (GAI) networks were higher than expected by chance (Figure 2b). While the three communities from the SRI network had distinct use of space, some communities from the GAI network had similar spatial distributions (e.g., communities yellow and purple, Figure 2c).

**Figure 2.**
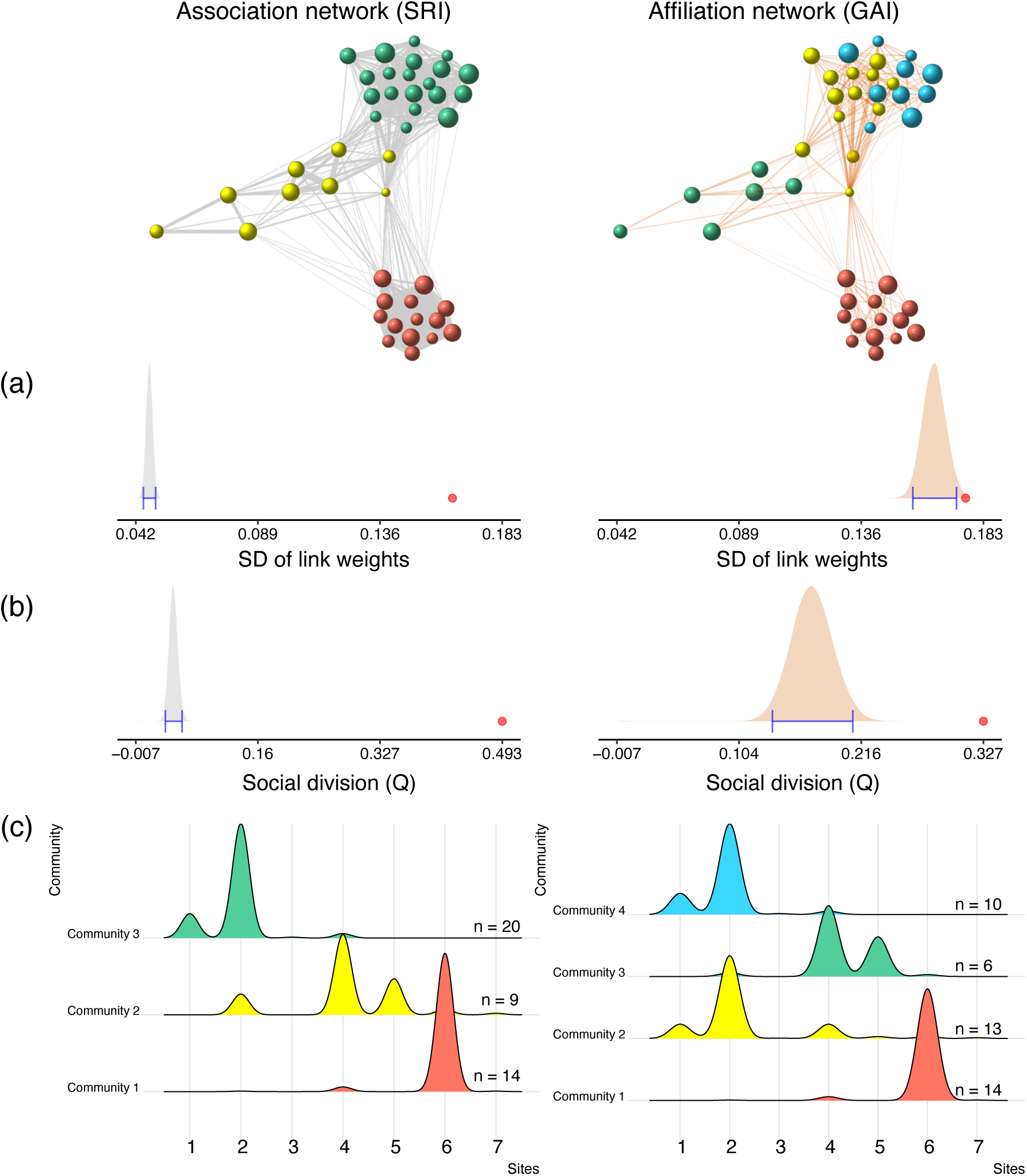
Shark social preferences at the individual and population levels. Nodes (N = 43; male:female = 30:13) representing photo-identified individuals are proportional to their size and colour-coded by social modules; individuals are connected by links whose thicknesses are proportional to SRI in the association networks (left panel), and to GAI removing confounding factors in the affiliation networks where only positive GAIs are plotted (right panel). In the density plots, red dots denote statistically significant observed values, shaded distributions indicate null expectancy and blue whiskers indicate 95% confidence intervals. The shark social network is characterized by (a) significant standard deviations (SD) of SRI and GAI indicating structured associations and social preferences, respectively, and (b) significant modularity (Q) indicating social division. In (c) are represented the density distribution of sightings of social community members.

### Crossing of genetic relatedness and association patterns

When testing for kin-biased relatedness, adult male-male (MM), female–female (FF) and male-female (MF) dyads did not have clear higher or lower genetic relatedness (Mantel test: MM, r = −0.012, n = 31, P = 0.061; FF, r = −0.019, n = 12, P = 0.685; MF, r = 0.023, n = 43, P = 0.259). In addition, mean genetic relatedness was not higher within than between size classes (mean r = 0.043, random 95% CI = 0.058-0.065, P = 0.88; Supplementary materials, Table S2). While individuals were relatively spatially clustered, genetic relatedness appeared much more homogeneously distributed across individuals and space (Mantel test between matrices of spatial overlap and genetic relatedness: r = 0.011, n = 43, P = 0.351; Supplementary Figure S4).

Average pairwise relatedness among individuals was 0.062 ± 0.001 (mean ± SE) ranging from 0 to 0.774. Associations were only significantly positively correlated with genetic relatedness between males (Mantel test: r= 0.103, P = 0.026) but no significant correlation was found between GAI and genetic relatedness for any sex dyad (Table 2). Males were generally more gregarious than females, as they significantly interacted with more individuals (higher degree) but did not have stronger relationships (higher strength) (Table 3, Figure 3). The lack of evidence that kinship drives associations in the shark network was also confirmed by a double premutation method (P_double permutation_ = 0.451).

**Table 2.** Results of Mantel test for the correlation between the simple-ratio association (SRI) and generalized affiliations (GAI) matrices and the pairwise genetic relatedness among individual sharks and each sex relationship. Observed estimates were compared to those of 20,000 estimates from the randomized networks and significant p-values are indicated in bold (proportion of times the empirical estimate was smaller than the null expectancy from 20,000 randomisations).

| Index | SRI |  | GAI |  |
| --- | --- | --- | --- | --- |
|  | Estimate (rho) | p-value | Estimate (rho) | p-value |
| Male-male (435 dyads) | 0.103 | **0.027** | 0.048 | 0.318 |
| Female-female (48 dyads) | -0.065 | 0.577 | 0.025 | 0.834 |
| Male-female (299 dyads) | 0.014 | 0.829 | -0.036 | 0.567 |

**Table 3.** Effects of sex on shark gregariousness (degree and strength) in the social network (N = 43; 30 males and 13 females). *P*-values (P_rand_) are given by the proportion of times the empirical regression coefficient was smaller than the null expectancy from 20,000 randomisations. P_double permutation_ is the p-value resulting from the double permutation procedure controlling for number of observations.

| | | Coefficient | SE | *t* statistic | $P_{rand}$ | $P_{double\ permutation}$ |
| --- | --- | --- | --- | --- | --- | --- |
| Degree | *Intercept* | 17.308 | 1.800 | 9.613 |  |  |
|  | Sex (male) | 4.726 | 2.156 | 2.192 | **< 0.001** | **0.031** |
| Strength | *Intercept* | 4.493 | 0.468 | 9.596 |  |  |
|  | Sex (male) | 0.369 | 0.560 | 0.660 | 0.548 | 0.513 |

**Figure 3.**
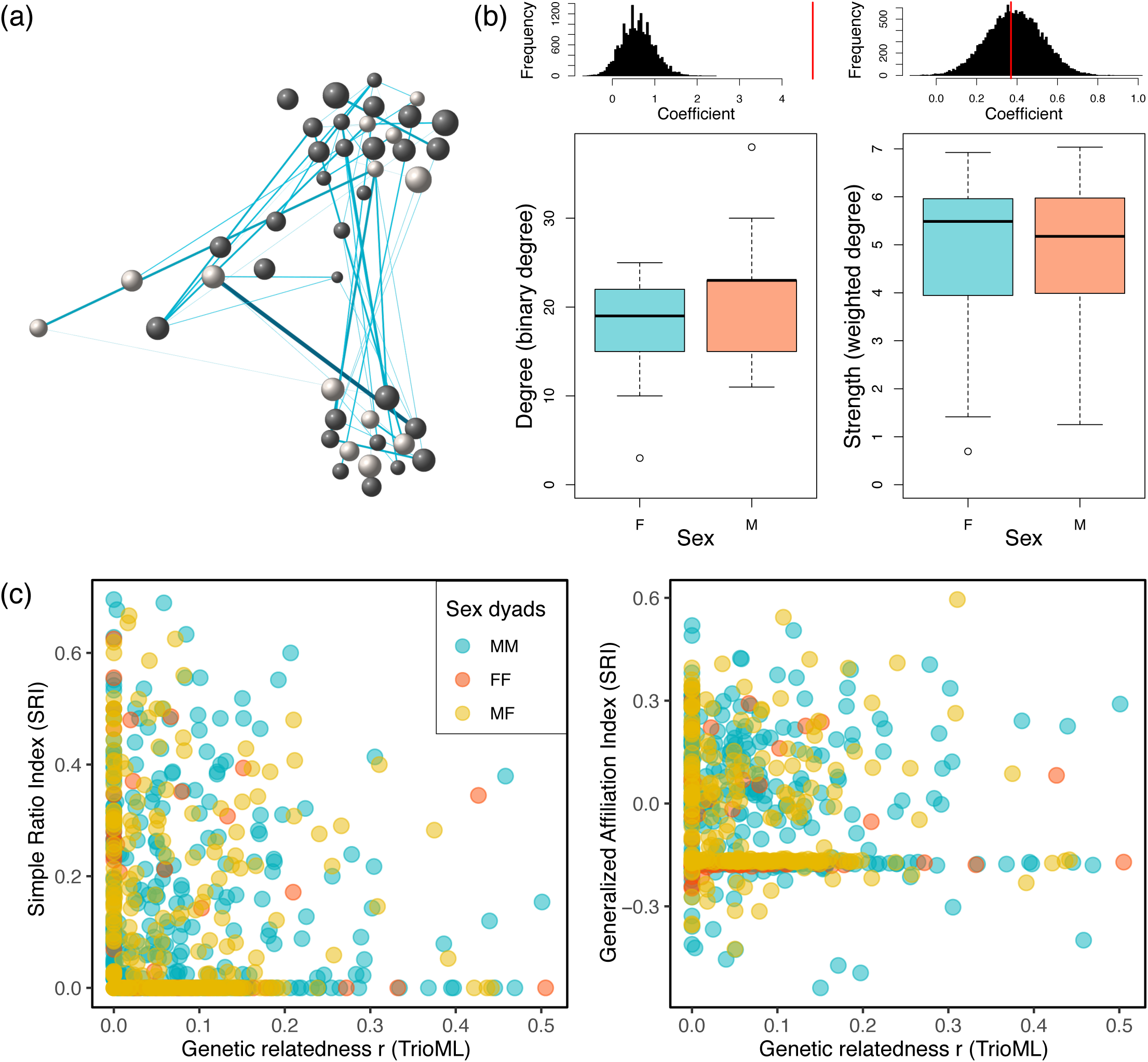
(a) Genetic network using the same layout as in Figure 2 where nodes (N = 43; male:female = 30:13) are proportional to individual size and colour-coded by sex (males in grey and females in white) and edges are proportional to genetic relatedness in the genetic network (only close kin are plotted, i.e. r >0.25). (b) Differences in gregariousness (degree and strength) of individual sharks between males and females (colours: pink indicates females and blue indicates males). Comparison of the coefficient from the GLM based on the observed data (red vertical line) and the frequency distribution of coefficients from the same model based on the randomized data are indicated over each box plot to report significance of the difference. (c) Relationship between association and affiliation indices and the triadic likelihood estimator (TrioML) of genetic relatedness in male-male, female-female and male-female pairs.

Within-community relatedness estimate was inferred for each community and index (SRI and GAI, Table 4). Relatedness within all communities was not higher than expected if communities were randomly organized (SRI network: within mean ± SE = 0.071 ± 0.005, between mean ± SE = 0.058 ± 0.003, P = 0.093; GAI network: within mean ± SE = 0.065 ± 0.006, between mean ± SE = 0.062 ± 0.006, P = 0.714) (Table 4).

**Table 4.** Community-level information on structure, association and affiliation indices, and genetic relatedness. For each social index (SRI and GAI) and each community, are reported number of community members (no. of individuals), mean index value, mean genetic relatedness and p-values of the one-tailed tests of within-group higher relatedness than random.

| Index | Communities | Modularity Q | No. of individuals | Mean index (SD) | Mean r (SE) | *p-values* |
| --- | --- | --- | --- | --- | --- | --- |
| SRI | Overall | 0.493 | 43 | 0.133 (0.210) | 0.071 (0.005) | 0.093 |
|  | Red |  | 14 | 0.487 (0.178) | 0.071 (0.009) | 0.267 |
|  | Yellow |  | 9 | 0.256 (0.307) | 0.073 (0.016) | 0.24 |
|  | Green |  | 20 | 0.282 (0.194) | 0.069 (0.006) | 0.11 |
| GAI | Overall | 0.312 | 43 | -0.028 (0.126) | 0.065 (0.006) | 0.552 |
|  | Red |  | 14 | 0.001 (0.148) | 0.072 (0.009) | 0.237 |
|  | Yellow |  | 13 | 0.085 (0.149) | 0.061 (0.009) | 0.652 |
|  | Green |  | 6 | 0.009 (0.152) | 0.043 (0.015) | 0.791 |
|  | Cyan |  | 10 | 0.047 (0.133) | 0.068 (0.015) | 0.235 |

Among the 903 potential pairs, 39 (4.31%) had genetic relatedness values higher than 0.25. In addition, there was no higher proportion of close relatives within than between communities for relatedness value r > 0.25 for SRI (chi-squared test: n_within/between_ = 17/ 22, d.f. = 1, χ^2^ = 0.928, P = 0.119) and GAI (chi-squared test: n_within/between_ = 11/29, d.f. = 1, χ^2^ = 0.017, P = 0.663) (Figure 4).

**Figure 4.**
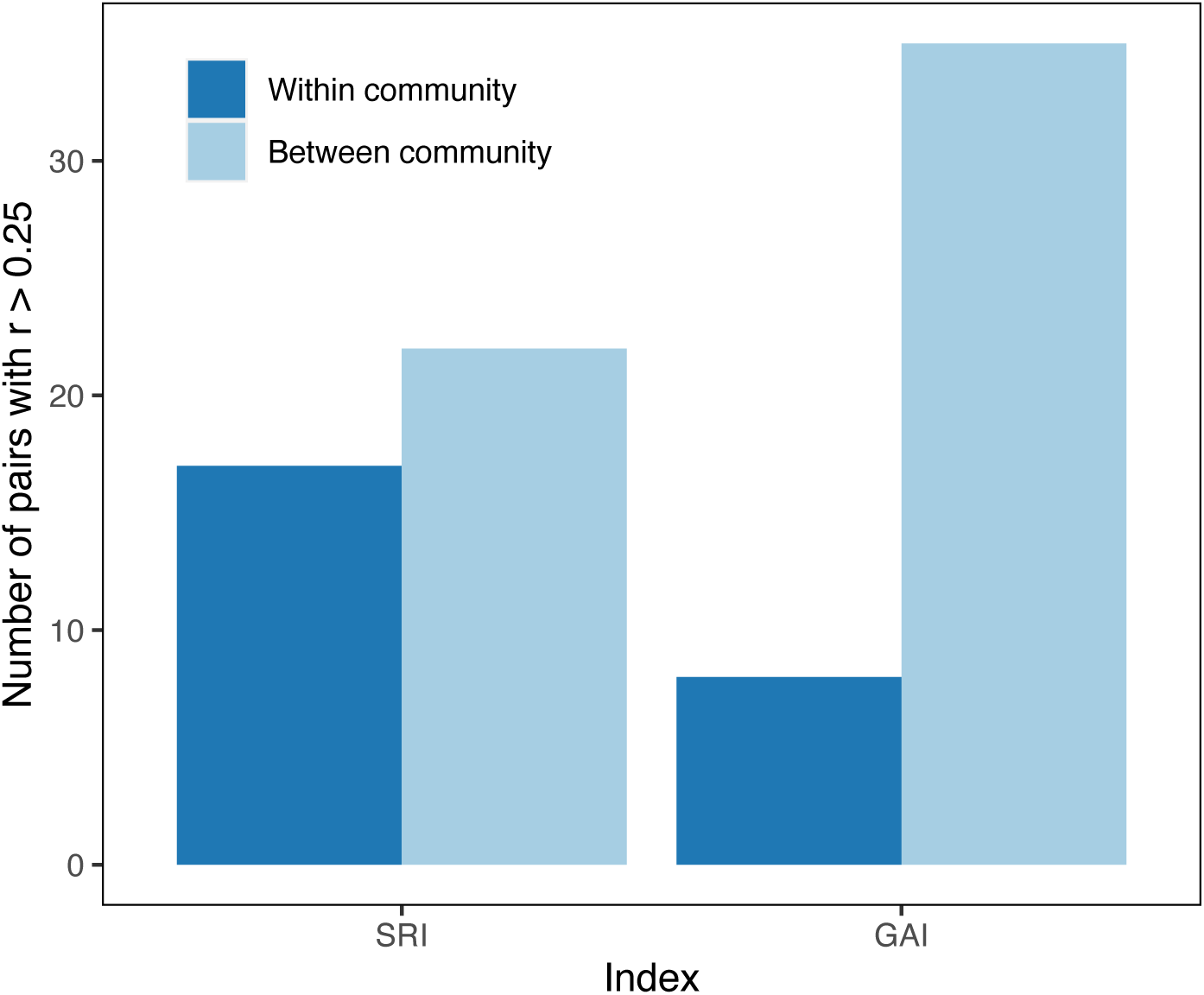
Numbers of close kin (r > 0.25, the expected value of half-sibs) within and between social communities for association (SRI) and affiliation (GAI) indices.

Together, these results suggest that no differences exist for within- and between-community membership with respect to the genetic relatedness of their members.

## DISCUSSION

Taking into account the confounding effects of 5 structural variables (spatial and temporal overlap, gregariousness, size and sex), which are known to influence association patterns (e.g. Godde et al. 2013; Diaz-Aguirre et al. 2019; Machado et al. 2019; Perryman et al. 2019), we found that blacktip reef sharks in Moorea had structured associations and affiliations, although most of the network structure was driven by spatiotemporal overlap. However, social proximity was not predicted by the genetic relatedness between sharks both at the association, affiliation and community levels. At the dyad level, only male-male associations, but not affiliations, were slightly correlated with genetic relatedness. In addition, individuals had low probabilities of interacting with a close kin which could explain the lack of influence of kinship in structuring the social network in this population. These results therefore suggest that genetic relatedness does not drive the structure of the social network in this shark population.

Compared to previous work conducted on this population (Mourier et al. 2012) which only analysed social structure through associations among sharks, the present study also considered the effects of two structural variables to estimate affiliation indices. Affiliations are an increasingly used method to investigate active social preferences experienced by animals (Whitehead and James 2015), in particular by considering the strong correlation that can exist between space use overlap and association indices in a variety of taxa (e.g., Mourier et al. 2012; Carter et al. 2013; Best et al. 2014). In our study, the network built from associations using the simple ratio index (SRI) was composed of three main communities relatively spatially separated and only low overlap (Figure 2). When removing the influence of spatial and temporal overlap from association patterns, the SD of the generalized affiliation indices (GAI) higher than random demonstrated the presence of active social preferences. The network built from the GAI revealed four communities that were less spatially separated, two of them having high spatial overlap and representing subcommunities spatially embedded in the communities detected by the network built from the SRI. This means that associations between sharks were the result of more than just similarities in habitat use or overlaps in time, indicating that individuals actively chose to group with preferred social partners. The differences between the three SRI communities in the present study and the four communities found in Mourier et al. (2012) can be due to the high threshold we used that may highlight only strong relationships and the use of SRI instead of HWI. To our knowledge, only one study on elasmobranchs has investigated social structure using GAI, demonstrating that manta rays also preferred affiliations (Perryman et al. 2019). Individuals’ site preferences and being present in the study at the same time was a strong predictor of association between pairs. Site fidelity is often a prerequisite for sociality, creating an environment for social relationships to develop and the emergence of social preferences. However, the presence of preferred social affiliations demonstrates that sharks show active social preferences that do not rely on preferences for sites and periods. Our study confirms that the observed shark social structure resembles that of a fission-fusion society characterized by an open and fluid social structure, long-term social recognition and a high number of potential affiliates, which is flexible depending on environmental conditions.

Among the adult sharks in our population, there was a generally low level of relatedness, and only a small number of dyads had close familial relatives. Interactions frequently occurred between distant kin and non-kin. This implies that the social structure among adult blacktip reef sharks was not based on associations between close kin as demonstrated by our analyses which compared association and affiliation patterns with genetic relatedness among dyads at the pairwise or community levels. This is confirmed by the low number of close kin available for each shark in the population (< 6 % pairs with r > 0.25), thereby limiting the probability of an individual to encounter a family member and to develop strong associations with them. The lack of genetic relatedness structure within size classes and the lack of decreasing genetic relatedness as sharks grow also suggests that juveniles are unlikely to leave their nursery ground with other kin. If young sharks were developing and maintaining strong bonds with their littermates throughout their entire life, we would have expected to find high mean relatedness and high proportion of close kin across all size classes. The low relatedness we found within each size class indicates that sharks favoured associations with non-kin. These results can be explained in part by the life history and life cycle of blacktip reef sharks. In fact, in contrast to most social animals that show some forms of family structure and parental care, female reef-associated sharks such as blacktip reef shark, leave their pups in their nursery after birth (Mourier and Planes 2013). Moreover, litter size in this species does not exceed five pups (Mourier, Mills, et al. 2013) while litter size in Moorea was limited to a maximum of two pups (Mourier and Planes 2013). In addition, blacktip reef sharks follow a yearly breeding cycle with females giving birth every year and potentially being fertilized by multiple males within or across years, which increases the probability of having maternal and paternal half-siblings. Our ongoing long-term nursery monitoring shows that capture probabilities rapidly decline after March (unpublished data), 2 to 3 months after parturition, which suggests a dramatic mortality rate within the nursery areas during the first months of life (i.e. survival rate expected to be inferior to 50% during the first year of life). Together with a small litter size and absence of parental care, this high mortality rate, which is common in many shark species, is likely to limit the opportunity to find family members and develop strong affiliations with close relatives at adulthood. Even in nurseries, juvenile lemon sharks did not clearly assort by relatedness (Guttridge et al. 2011), even if the probability of finding a relative is higher for this species with a larger litter size. When juvenile sharks grow, they progressively explore their environment and increase their home range (Chin A et al. 2013), creating an opportunity to find related individuals such as parents or maternal half-siblings from previous reproductive seasons. At adulthood, our results confirm that preferred associations and affiliations are not driven by genetic relatedness as sharks are associating with conspecifics of variable genetic distances. This suggests that sharks might not have the ability for kin recognition simply based on visual or olfactory cues and that kin-based preferred associations and affiliations may only develop within nursery areas from increased familiarity with littermates, or that they are not seeking for associations with related individuals. Through investigation of social groups of spotted eagle rays *Aetobatus narinari* in Florida, Newby et al. (2014) found no kin-structure in the social organization, although the analysis was based on group composition rather that quantitatively inferred using association indices. However, our results revealed that males slightly preferred to associate with other related males but this tendency was not confirmed for affiliations (accounting for spatial and temporal structural components). This suggests that genetic relatedness among males was spatially structured and that males may disperse less than females. The lack of differences in relatedness between males and females suggests that the risk of inbreeding might be low if these interactions represented potential mating pairs and not only social bonds.

The emerging literature suggests that genetic structure of animal social networks can vary dramatically, from highly cohesive kin‐based groups like African elephants (*Loxodonta africana*) (Archie et al. 2006) or spotted hyenas (*Crocuta crocuta*) (Holekamp et al. 2012), to groups with moderate levels of genetic relatedness due to limited dispersal like the Galápagos sea lion (*Zalophus wollebaeki*) (Wolf and Trillmich 2008) or the eastern grey kangaroo (*Macropus giganteus*) (Best et al. 2014), or to groups with little to no genetic relatedness like guppies (*Poecilia reticulata*) (Croft et al. 2012), the common raccoon (*Procyon lotor*) (Hirsch et al. 2013) or migratory golden‐crowned sparrows (*Zonotrichia atricapilla*) (Arnberg et al. 2015). These patterns of variation provide opportunities to explore how ecological factors interact with kinship to produce variations in the structures of animal societies. Kinship is expected to promote the evolution of cooperation and sociality in animals (Hamilton 1964). However, our understanding of the evolution of sociality results to a great extent from the study of closed societies, in which interactions mainly involve relatives and can hence be explained by kin selection (Hamilton 1964). However, the kin selection theory has recently been challenged by results from studies showing that fitness benefit can emerge in social groups composed mainly of non-relatives (e.g., Cameron et al. 2009; Riehl 2011; Wilkinson et al. 2016). In many natural populations, dispersal tends to be limited, favouring local competition between neighbours and the emergence of a social component, whether it be selfish, aggressive, cooperative or altruistic (Lehmann and Rousset 2010). But how social behaviours translate into fitness costs and benefits depends considerably on life-history features, as well as on local demographic and ecological conditions. The fission – fusion social dynamics lead to unstable group membership, and dispersal and occasional recruitment of unrelated individuals lead to low average relatedness in groups. Then under such conditions, selection is not expected to favour kin recognition mechanisms based on familiarity alone.

Therefore, contrary to the kin selection hypothesis which predicts stronger associations among kin, sharks tended to assort randomly according to relatedness. As kinship does not explain the strength of social affiliations in blacktip reef sharks, the question remains as to how and why sharks form preferred associations and affiliations organised in social communities (Mourier et al. 2012). Although cooperation has been mainly explained in the context of kin selection, there might be potential benefits of non-kin sociality in blacktip reef sharks such as for other animals in which association with non-kin emerges via reciprocal altruism (Carter and Wilkinson 2013; Wilkinson et al. 2016). In addition, by-product mutualism and pseudo-reciprocity are simple mechanisms that can lead to increased foraging success, cooperative hunting. While evidence of shark cooperation has not been confirmed, gregarious behaviour can have several benefits in sharks (Jacoby et al. 2012), including increased foraging success by hunting in groups (Weideli et al. 2015; Mourier et al. 2016), protection from predators (Mourier, Planes, et al. 2013), or increased tolerance relationships and reduced aggression rate (Brena et al. 2018). Heterospecific foraging associations have been found to develop and increase predation success (Labourgade et al. 2020), which suggests that sharks can benefit from hunting associations without associating with kin. These benefits do not necessarily imply kin selection and can simply build on the development of familiarity from repeated interactions. Social structure in reef sharks can arise from multiple simple ecological factors such as the distribution of resources in space and time leading to aggregations of individuals even in the absence of benefits of direct social affiliation (Ramos-Fernández et al. 2006) or mitigation of the cost of unnecessary aggression when competing for resources mediated by individual recognition (Brena et al. 2018). Regardless of the exact cause of social preferences in reef sharks, the absence of kinship as an important factor in structuring association patterns suggests that there are important benefits of sociality in sharks that we still need to uncover. With an increasing use of social network analyses applied to shark populations (Mourier et al. 2018), future work on social networks and genetic relatedness in different populations or species is necessary to confirm our results and to improve our understanding of population dynamics in sharks and the evolution of sociality.

## ACKNOWLEDGEMENTS

We are grateful to the Centre de Recherche Insulaire et Observatoire de l’Environnement (CRIOBE) staff, volunteers and students who assisted in field data collection. We are also grateful to Jeanine Almany for English reviewing.

## ETHICS STATMENT

This work was approved by the Direction à l’Environnement (DIREN) of French Polynesia and Ministère de la promotion des langues, de la culture, de la communication et de l’environnement de Polynésie française under Arrêté 9324 of 30 October 2015.

### Data accessibility

Data are available on dryad: to be deposited if accepted.

## SUPPLEMENTARY MATERIAL

Supplementary information including two tables and four figures can be found online.

## REFERENCES

Archie EA, Moss CJ, Alberts SC. 2006. The ties that bind: genetic relatedness predicts the fission and fusion of social groups in wild African elephants. Proc R Soc Lond B Biol Sci. 273(1586):513–522. doi: 10.1098/rspb.2005.3361.

Arnberg NN, Shizuka D, Chaine AS, Lyon BE. 2015. Social network structure in wintering golden□crowned sparrows is not correlated with kinship. Mol Ecol. 24(19):5034–5044.

Bejder L, Fletcher D, Bräger S. 1998. A method for testing association patterns of social animals. Anim Behav. 56(3):719–725. doi: 10.1006/anbe.1998.0802.

Best EC, Dwyer RG, Seddon JM, Goldizen AW. 2014. Associations are more strongly correlated with space use than kinship in female eastern grey kangaroos. Anim Behav. 89:1–10. doi: 10.1016/j.anbehav.2013.12.011.

Brena PF, Mourier J, Planes S, Clua EE. 2018. Concede or clash? Solitary sharks competing for food assess rivals to decide. Proc R Soc B. 285(1875):20180006. doi: 10.1098/rspb.2018.0006.

Brown GE, Brown JA. 1993. Social dynamics in salmonid fishes: do kin make better neighbours? Anim Behav. 45(5):863–871. doi: 10.1006/anbe.1993.1107.

Buston PM, Fauvelot C, Wong MYL, Planes S. 2009. Genetic relatedness in groups of the humbug damselfish Dascyllus aruanus: small, similar-sized individuals may be close kin. Mol Ecol. 18(22):4707–4715. doi: 10.1111/j.1365-294X.2009.04383.x.

Cairns SJ, Schwager SJ. 1987. A comparison of association indices. Anim Behav. 35(5):1454–1469. doi: 10.1016/S0003-3472(87)80018-0.

Cameron EZ, Setsaas TH, Linklater WL. 2009. Social bonds between unrelated females increase reproductive success in feral horses. Proc Natl Acad Sci. 106(33):13850–13853. doi: 10.1073/pnas.0900639106.

Carter GG, Wilkinson GS. 2013. Food sharing in vampire bats: reciprocal help predicts donations more than relatedness or harassment. Proc R Soc B Biol Sci. 280(1753):20122573. doi: 10.1098/rspb.2012.2573.

Carter KD, Seddon JM, Frère CH, Carter JK, Goldizen AW. 2013. Fission–fusion dynamics in wild giraffes may be driven by kinship, spatial overlap and individual social preferences. Anim Behav. 85(2):385–394. doi: 10.1016/j.anbehav.2012.11.011.

Chapman DD, Babcock EA, Gruber SH, Dibattista JD, Franks BR, Kessel SA, Guttridge T, Pikitch EK, Feldheim KA. 2009. Long-term natal site-fidelity by immature lemon sharks (Negaprion brevirostris) at a subtropical island. Mol Ecol. 18(16):3500–3507. doi: 10.1111/j.1365-294X.2009.04289.x.

Chin A, Heupel MR, Simpfendorfer CA, Tobin AJ. 2013. Ontogenetic movements of juvenile blacktip reef sharks: evidence of dispersal and connectivity between coastal habitats and coral reefs. Aquat Conserv Mar Freshw Ecosyst. 23(3):468–474. doi: 10.1002/aqc.2349.

Clutton-Brock T. 2009. Cooperation between non-kin in animal societies. Nature. 462(7269):51–57. doi: 10.1038/nature08366.

Croft DP, Hamilton PB, Darden SK, Jacoby DMP, James R, Bettaney EM, Tyler CR. 2012. The role of relatedness in structuring the social network of a wild guppy population. Oecologia. 170(4):955–963. doi: 10.1007/s00442-012-2379-8.

Dadda M, Pilastro A, Bisazza A. 2005. Male sexual harassment and female schooling behaviour in the eastern mosquitofish. Anim Behav. 70(2):463–471. doi: 10.1016/j.anbehav.2004.12.010.

Diaz-Aguirre F, Parra GJ, Passadore C, Möller L. 2019. Genetic relatedness delineates the social structure of southern Australian bottlenose dolphins. Behav Ecol. 30(4):948–959. doi: 10.1093/beheco/arz033.

Dixon P. 2003. VEGAN, a package of R functions for community ecology. J Veg Sci. 14(6):927–930. doi: 10.1111/j.1654-1103.2003.tb02228.x.

Farine DR. 2013. Animal social network inference and permutations for ecologists in R using asnipe. Methods Ecol Evol. 4(12):1187–1194. doi: 10.1111/2041-210X.12121.

Farine DR. 2017. A guide to null models for animal social network analysis. Methods Ecol Evol.:n/a-n/a. doi: 10.1111/2041-210X.12772.

Farine DR, Carter GG. 2020 Aug 4. Permutation tests for hypothesis testing with animal social data: problems and potential solutions. bioRxiv.:2020.08.02.232710. doi: 10.1101/2020.08.02.232710.

Farine DR, Whitehead H. 2015. Constructing, conducting and interpreting animal social network analysis. J Anim Ecol. 84(5):1144–1163. doi: 10.1111/1365-2656.12418.

Feldheim KA, Gruber SH, DiBattista JD, Babcock EA, Kessel ST, Hendry AP, Pikitch EK, Ashley MV, Chapman DD. 2014. Two decades of genetic profiling yields first evidence of natal philopatry and long□term fidelity to parturition sites in sharks. Mol Ecol. 23(1):110–117. doi: 10.1111/mec.12583.

Franks DW, Ruxton GD, James R. 2010. Sampling Animal Association Networks with the Gambit of the Group. Behav Ecol Sociobiol. 64(3):493–503.

Franks DW, Weiss MN, Silk MJ, Perryman RJY, Croft DP. 2020. Calculating effect sizes in animal social network analysis. Methods Ecol Evol. n/a(n/a). doi: 10.1111/2041-210X.13429. [accessed 2020 Sep 28]. https://besjournals.onlinelibrary.wiley.com/doi/abs/10.1111/2041-210X.13429.

Godde S, Humbert L, Côté SD, Réale D, Whitehead H. 2013. Correcting for the impact of gregariousness in social network analyses. Anim Behav. 85(3):553–558. doi: 10.1016/j.anbehav.2012.12.010.

Guttridge TL, Dijk S van, Stamhuis EJ, Krause J, Gruber SH, Brown C. 2013. Social learning in juvenile lemon sharks, Negaprion brevirostris. Anim Cogn. 16(1):55–64. doi: 10.1007/s10071-012-0550-6.

Guttridge TL, Gruber SH, DiBattista JD, Feldheim KA, Croft DP, Krause S, Krause J. 2011. Assortative interactions and leadership in a free-ranging population of juvenile lemon shark Negaprion brevirostris. Mar Ecol Prog Ser. 423:235–245. doi: 10.3354/meps08929.

Guttridge TL, Gruber SH, Gledhill KS, Croft DP, Sims DW, Krause J. 2009. Social preferences of juvenile lemon sharks, Negaprion brevirostris. Anim Behav. 78(2):543–548. doi: 10.1016/j.anbehav.2009.06.009.

Hamilton WD. 1964. The genetical evolution of social behaviour. I. J Theor Biol. 7(1):1–16. doi: 10.1016/0022-5193(64)90038-4.

Hatchwell Ben J. 2010. Cryptic Kin Selection: Kin Structure in Vertebrate Populations and Opportunities for Kin□Directed Cooperation. Ethology. 116(3):203–216. doi: 10.1111/j.1439-0310.2009.01732.x.

Hirsch BT, Prange S, Hauver SA, Gehrt SD. 2013. Genetic relatedness does not predict racoon social network structure. Anim Behav. 85(2):463–470. doi: 10.1016/j.anbehav.2012.12.011.

Holekamp KE, Smith JE, Strelioff CC, Van Horn RC, Watts HE. 2012. Society, demography and genetic structure in the spotted hyena. Mol Ecol. 21(3):613–632. doi: 10.1111/j.1365-294X.2011.05240.x.

Hoppitt WJE, Farine DR. 2018. Association indices for quantifying social relationships: how to deal with missing observations of individuals or groups. Anim Behav. 136:227–238. doi: 10.1016/j.anbehav.2017.08.029.

Jacoby DMP, Busawon DS, Sims DW. 2010. Sex and social networking: the influence of male presence on social structure of female shark groups. Behav Ecol. 21(4). doi: 10.1093/beheco/arq061. [accessed 2017 Jan 4]. http://beheco.oxfordjournals.org/content/early/2010/05/09/beheco.arq061.

Jacoby DMP, Croft DP, Sims DW. 2012. Social behaviour in sharks and rays: analysis, patterns and implications for conservation. Fish Fish. 13(4):399–417. doi: 10.1111/j.1467-2979.2011.00436.x.

Jacoby DMP, Papastamatiou YP, Freeman R. 2016. Inferring animal social networks and leadership: applications for passive monitoring arrays. J R Soc Interface. 13(124):20160676. doi: 10.1098/rsif.2016.0676.

Kerth G, Perony N, Schweitzer F. 2011. Bats are able to maintain long-term social relationships despite the high fission–fusion dynamics of their groups. Proc R Soc Lond B Biol Sci. 278(1719):2761–2767. doi: 10.1098/rspb.2010.2718.

Krause J, Ruxton GD. 2002. Living in Groups. OUP Oxford.

Labourgade P, Ballesta L, Huveneers C, Papastamatiou Y, Mourier J. 2020. Heterospecific foraging associations between reef-associated sharks: first evidence of kleptoparasitism in sharks. Ecology.

Landeau L, Terborgh J. 1986. Oddity and the ‘confusion effect’ in predation. Anim Behav. 34(5):1372–1380. doi: 10.1016/S0003-3472(86)80208-1.

Lehmann L, Rousset F. 2010. How life history and demography promote or inhibit the evolution of helping behaviours. Philos Trans R Soc Lond B Biol Sci. 365(1553):2599–2617. doi: 10.1098/rstb.2010.0138.

Li CC, Weeks DE, Chakravarti A. 1993. Similarity of DNA fingerprints due to chance and relatedness. Hum Hered. 43(1):45–52.

Lukas D, Reynolds V, Boesch C, Vigilant L. 2005. To what extent does living in a group mean living with kin? Mol Ecol. 14(7):2181–2196. doi: 10.1111/j.1365-294X.2005.02560.x.

Lynch M, Ritland K. 1999. Estimation of Pairwise Relatedness With Molecular Markers. Genetics. 152:1753–1766.

Machado AMS, Cantor M, Costa APB, Righetti BPH, Bezamat C, Valle-Pereira JVS, Simões-Lopes PC, Castilho PV, Daura-Jorge FG. 2019. Homophily around specialized foraging underlies dolphin social preferences. Biol Lett. 15(4):20180909. doi: 10.1098/rsbl.2018.0909.

Mann J, Stanton MA, Patterson EM, Bienenstock EJ, Singh LO. 2012. Social networks reveal cultural behaviour in tool-using dolphins. Nat Commun. 3:980. doi: 10.1038/ncomms1983.

Milligan BG. 2003. Maximum-Likelihood Estimation of Relatedness. Genetics. 163(3):1153–1167.

Mourier J, Brown C, Planes S. 2017. Learning and robustness to catch-and-release fishing in a shark social network. Biol Lett. 13(3):20160824. doi: 10.1098/rsbl.2016.0824.

Mourier J, Lédée EJI, Guttridge TL, Jacoby DMP. 2018. Network analysis and theory in shark ecology - methods and applications. In: Shark Research: Emerging Technologies and Application for the Study of Shark Biology. Carrier J, Heithaus M and Simpfendorfer C. CRC Press. p. 337–356.

Mourier J, Maynard J, Parravicini V, Ballesta L, Clua E, Domeier ML, Planes S. 2016. Extreme Inverted Trophic Pyramid of Reef Sharks Supported by Spawning Groupers. Curr Biol. 26(15):2011–2016. doi: 10.1016/j.cub.2016.05.058.

Mourier J, Mills SC, Planes S. 2013. Population structure, spatial distribution and life-history traits of blacktip reef sharks Carcharhinus melanopterus. J Fish Biol. 82(3):979–993. doi: 10.1111/jfb.12039.

Mourier J, Planes S. 2013. Direct genetic evidence for reproductive philopatry and associated fine-scale migrations in female blacktip reef sharks (Carcharhinus melanopterus) in French Polynesia. Mol Ecol. 22(1):201–214. doi: 10.1111/mec.12103.

Mourier J, Planes S, Buray N. 2013. Trophic interactions at the top of the coral reef food chain. Coral Reefs. 32(1):285–285. doi: 10.1007/s00338-012-0976-y.

Mourier J, Vercelloni J, Planes S. 2012. Evidence of social communities in a spatially structured network of a free-ranging shark species. Anim Behav. 83(2):389–401. doi: 10.1016/j.anbehav.2011.11.008.

Newby J, Darden T, Bassos-Hull K, Shedlock AM. 2014. Kin structure and social organization in the spotted eagle ray, Aetobatus narinari, off coastal Sarasota, FL. Environ Biol Fishes. 97(9):1057–1065. doi: 10.1007/s10641-014-0289-9.

Newman MEJ. 2006. Finding community structure in networks using the eigenvectors of matrices. Phys Rev E. 74(3):036104. doi: 10.1103/PhysRevE.74.036104.

Olsén KH, JäUrvi T. 2005. Effects of kinship on aggression and RNA content in juvenile Arctic charr. J Fish Biol. 51(2):422–435. doi: 10.1111/j.1095-8649.1997.tb01676.x.

Papastamatiou YP, Lowe CG, Caselle JE, Friedlander AM. 2009. Scale-dependent effects of habitat on movements and path structure of reef sharks at a predator-dominated atoll. Ecology. 90(4):996–1008. doi: 10.1890/08-0491.1.

Perryman RJY, Venables SK, Tapilatu RF, Marshall AD, Brown C, Franks DW. 2019. Social preferences and network structure in a population of reef manta rays. Behav Ecol Sociobiol. 73(8):114. doi: 10.1007/s00265-019-2720-x.

Pew J, Muir PH, Wang J, Frasier TR. 2015. related: an R package for analysing pairwise relatedness from codominant molecular markers. Mol Ecol Resour. 15(3):557–561. doi: 10.1111/1755-0998.12323.

Puga□Gonzalez I, Sueur C, Sosa S. 2020. Null models for animal social network analysis and data collected via focal sampling: Pre-network or node network permutation? Methods Ecol Evol. n/a(n/a). doi: 10.1111/2041-210X.13400. [accessed 2020 Sep 28]. https://besjournals.onlinelibrary.wiley.com/doi/abs/10.1111/2041-210X.13400.

Queller DC, Goodnight KF. 1989. Estimating Relatedness Using Genetic Markers. Evolution. 43(2):258–275. doi: 10.1111/j.1558-5646.1989.tb04226.x.

R Core Team. 2015. R: a language and environment for statistical computing. Vienna, Austria: R Computing Foundation for Science.

Ramos-Fernández G, Boyer D, Gómez VP. 2006. A complex social structure with fission–fusion properties can emerge from a simple foraging model. Behav Ecol Sociobiol. 60(4):536–549. doi: 10.1007/s00265-006-0197-x.

Reisinger RR, Beukes (née Janse van Rensburg) C, Hoelzel AR, de Bruyn PJN. 2017. Kinship and association in a highly social apex predator population, killer whales at Marion Island. Behav Ecol. 28(3):750–759. doi: 10.1093/beheco/arx034.

Riehl C. 2011. Living with strangers: direct benefits favour non-kin cooperation in a communally nesting bird. Proc R Soc Lond B Biol Sci. 278(1712):1728–1735. doi: 10.1098/rspb.2010.1752.

Ritland K. 1996. Estimators for pairwise relatedness and individual inbreeding coefficients. Genet Res. 67(2):175–185. doi: 10.1017/S0016672300033620.

Snijders L, Blumstein DT, Stanley CR, Franks DW. 2017. Animal Social Network Theory Can Help Wildlife Conservation. Trends Ecol Evol. 32(8):567–577. doi: 10.1016/j.tree.2017.05.005.

Van Oosterhout Cock, Hutchinson William F., Wills Derek P. M., Shipley Peter. 2004. micro□checker: software for identifying and correcting genotyping errors in microsatellite data. Mol Ecol Notes. 4(3):535–538. doi: 10.1111/j.1471-8286.2004.00684.x.

Vignaud T, Clua E, Mourier J, Maynard J, Planes S. 2013. Microsatellite Analyses of Blacktip Reef Sharks (Carcharhinus melanopterus) in a Fragmented Environment Show Structured Clusters. PLOS ONE. 8(4):e61067. doi: 10.1371/journal.pone.0061067.

Vignaud TM, Mourier J, Maynard JA, Leblois R, Spaet JLY, Clua E, Neglia V, Planes S. 2014. Blacktip reef sharks, Carcharhinus melanopterus, have high genetic structure and varying demographic histories in their Indo-Pacific range. Mol Ecol. 23(21):5193–5207. doi: 10.1111/mec.12936.

Wang J. 2002. An Estimator for Pairwise Relatedness Using Molecular Markers. Genetics. 160(3):1203–1215.

Wang J. 2007. Triadic IBD coefficients and applications to estimating pairwise relatedness. Genet Res. 89(3):135–153. doi: 10.1017/S0016672307008798.

Weideli OC, Mourier J, Planes S. 2015. A massive surgeonfish aggregation creates a unique opportunity for reef sharks. Coral Reefs. 34(3):835–835. doi: 10.1007/s00338-015-1290-2.

Weiss MN, Franks DW, Brent LJN, Ellis S, Silk MJ, Croft DP. 2020 Apr 30. Common datastream permutations of animal social network data are not appropriate for hypothesis testing using regression models. bioRxiv.:2020.04.29.068056. doi: 10.1101/2020.04.29.068056.

Whitehead H. 2008. Analyzing Animal Societies: Quantitative Methods for Vertebrate Social Analysis. Chicago: University of Chicago Press.

Whitehead H, Dufault S. 1999. Techniques for Analyzing Vertebrate Social Structure Using Identified Individuals: Review and Recommendations. In: Slater PJB, Rosenblat JS, Snowden CT, Roper TJ, editors. Advances in the Study of Behavior. Vol. 28. Academic Press. p. 33–74. [accessed 2020 Apr 20]. http://www.sciencedirect.com/science/article/pii/S0065345408602156.

Whitehead H, James R. 2015. Generalized affiliation indices extract affiliations from social network data. Methods Ecol Evol. 6(7):836–844. doi: 10.1111/2041-210X.12383.

Wilkinson GS, Carter GG, Bohn KM, Adams DM. 2016. Non-kin cooperation in bats. Phil Trans R Soc B. 371(1687):20150095. doi: 10.1098/rstb.2015.0095.

Wiszniewski J, Lusseau D, Möller LM. 2010. Female bisexual kinship ties maintain social cohesion in a dolphin network. Anim Behav. 80(5):895–904. doi: 10.1016/j.anbehav.2010.08.013.

Wittemyer G, Okello JBA, Rasmussen HB, Arctander P, Nyakaana S, Douglas-Hamilton I, Siegismund HR. 2009. Where sociality and relatedness diverge: the genetic basis for hierarchical social organization in African elephants. Proc R Soc B Biol Sci. 276(1672):3513–3521. doi: 10.1098/rspb.2009.0941.

Wolf JBW, Trillmich F. 2008. Kin in space: social viscosity in a spatially and genetically substructured network. Proc R Soc Lond B Biol Sci. 275(1647):2063–2069. doi: 10.1098/rspb.2008.0356.

