## Supplementary for "Kinship does not predict the structure of a shark social network" (from another version of this preprint: version 2, 2020-05-19)

### Supplementary materials

**Figure S1** Density plots of number of sightings per individuals. Dotted line represents the mean and dashed line represents the median.

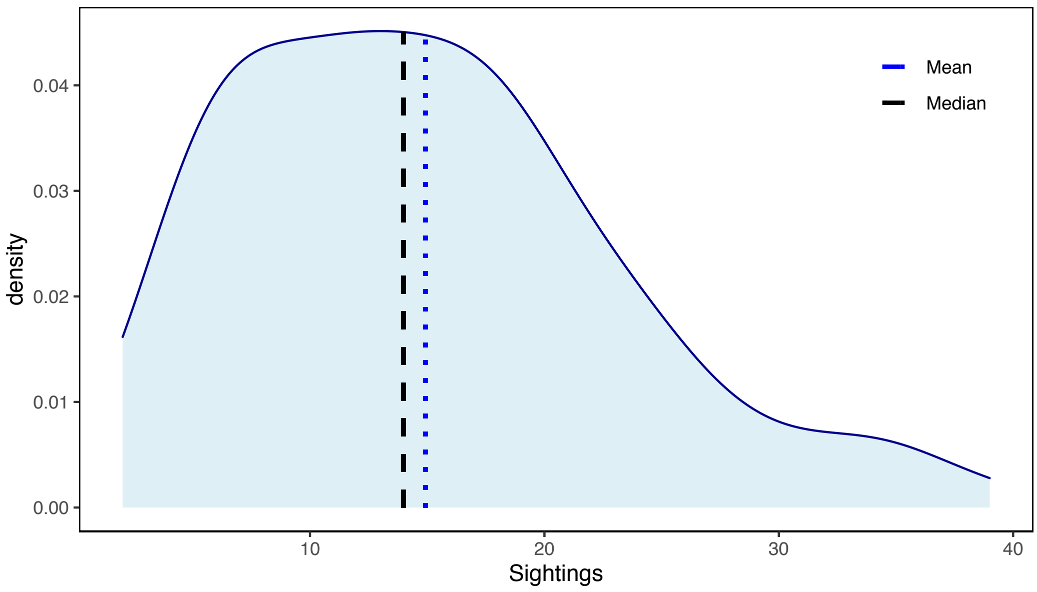

**Figure S2** Box plot comparing the relatedness estimates for pairs of simulated individuals of known relatedness using the best four of seven estimators tested. The estimators are coded as follows: dyadML (Milligan 2003), Q & G  (Queller and Goodnight 1989), trioML (Wang 2007) and W (Wang 2002). The standard deviations of estimators are indicated in blue, with the smallest SD in bold. Correlations with true values were: dyadML = 0.839, Q & G = 0.810, trioML = 0.840, W = 0.815.

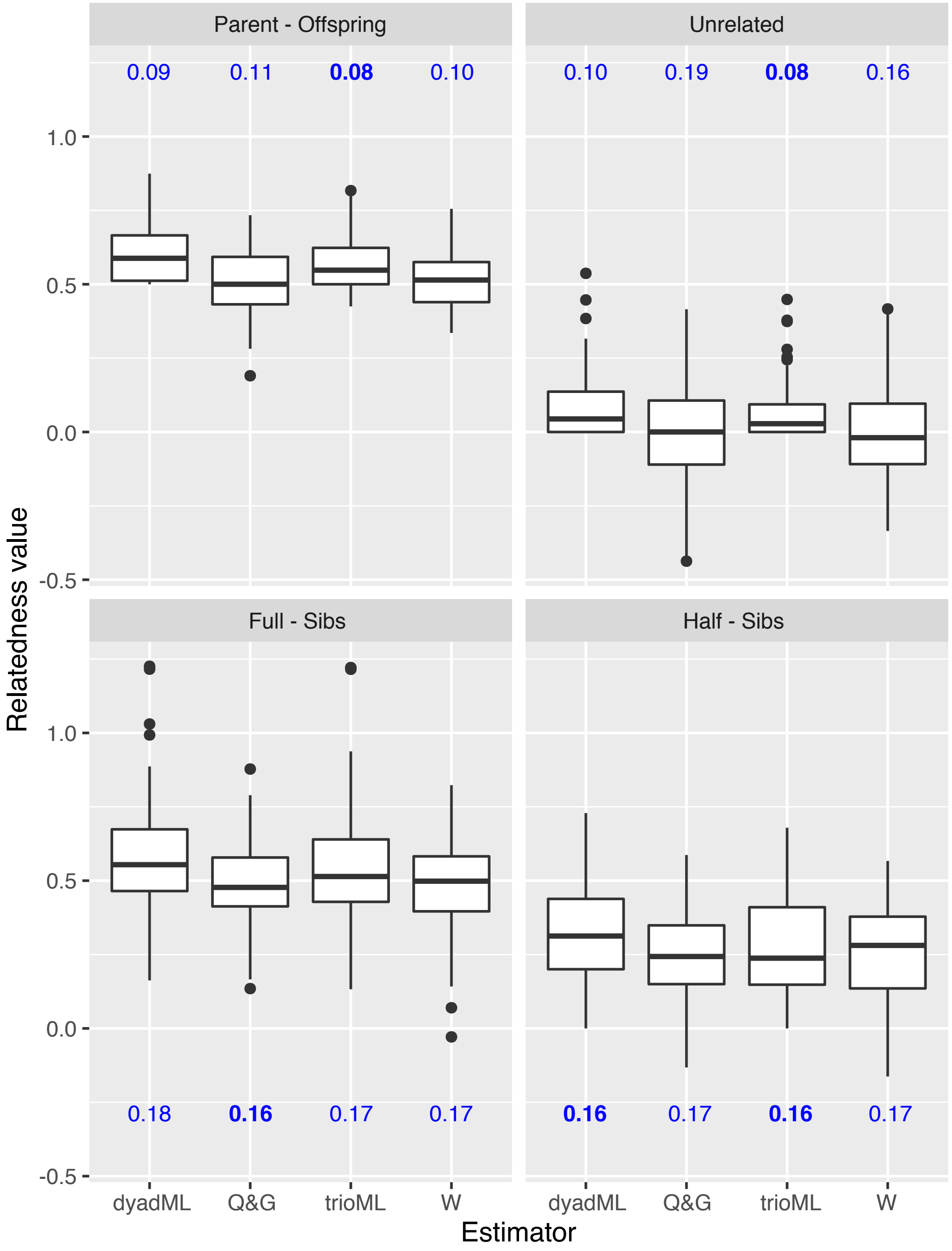

**Table S1** Summary statistics of heterozygosity and FIS of the 16 loci used in this study for 255 adult blacktip reef sharks. More details on microsatellite markers can be found in Mourier and Planes 2013.

| Heterozygosity | | | $F_{IS}$ | |
|---|---|---|---|---|
| Ho | He | P | $F_{IS}$ | P |
| 0.5888 | 0.5840 | 1.000 | -0.00603 | 0.298 |

Table S2: Mean observed within size classes relatedness. The expected distribution of within-community relatedness values if communities were randomly organized at their original sample size and based on 1000 iterations is indicated by mean and SD of random. P-values indicate the percentage of randomized iterations where the expected values were greater than or equal to the observed value.

| Size classes | N | mean r values | Mean (SD) of random | p-values |
|---|---|---|---|---|
| <110 | 1 | / | / | / |
| 110-119 | 14 | 0.053 (0.074) | 0.062 (0.016) | 0.73 |
| 120-129 | 8 | 0.040 (0.053) | 0.064 (0.023) | 0.89 |
| 130-139 | 13 | 0.045 (0.078) | 0.062 (0.015) | 0.87 |
| 140-149 | 4 | 0.050 (0.061) | 0.057 (0.038) | 0.56 |
| >150 | 3 | 0.034 (0.059) | 0.062 (0.063) | 0.54 |
| Total | 43 | 0.048 (0.073) | 0.061 (0.018) | 0.88 |

**Figure S3:** Spatial profiles of individual sharks represented by heatmap. Each row corresponds to an individual and cell color is proportional to its encounter rate (i.e., no. sightings of individual at site, divided by no. sampling occasions at site) at each site (from sites 1 to 7, Figure 1) on each column. Empty color (i.e. white) cells indicate that no sighting was recorded at the corresponding site.

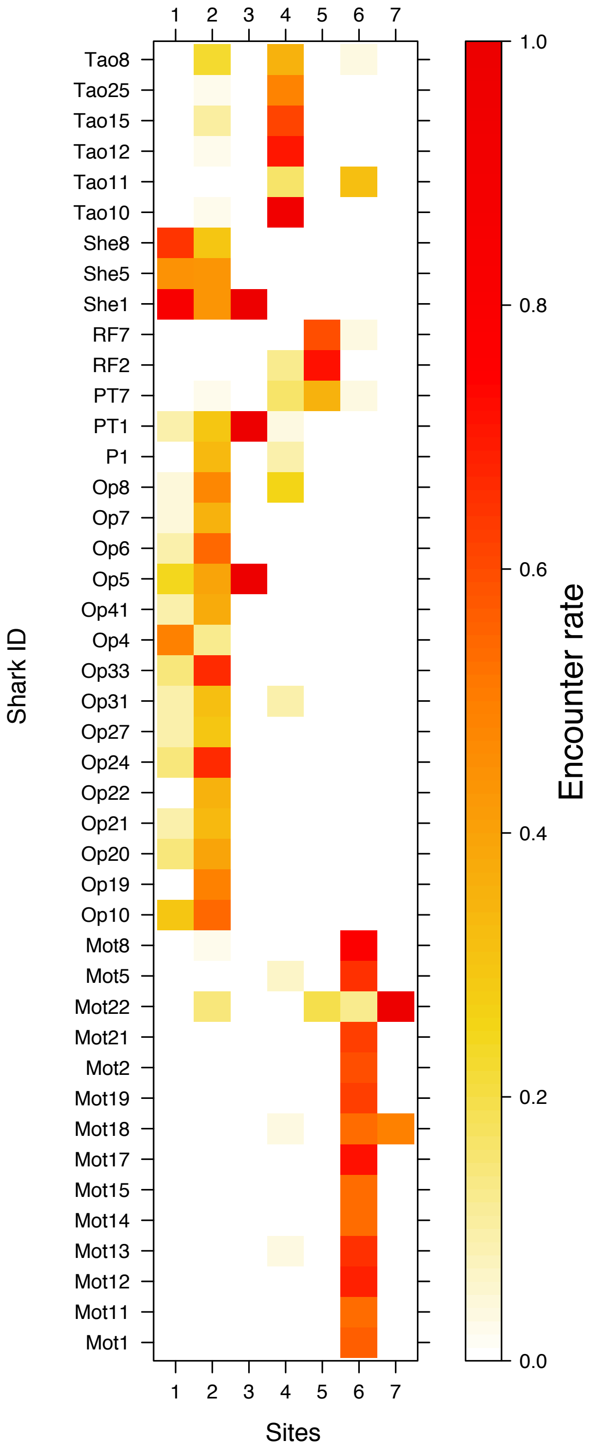

**Figure S4:** The subset of 42 individual sharks for which all individual traits were available: size and sex classes, spatial overlap, temporal overlap, and genetic relatedness. Alphanumeric codes indicate individual photo-identification labels along with sex. In the upper triangle, the colour code indicates spatial overlap (SO) between pairs of individuals. In the lower triangle, the colour code indicates genetic relatedness between individuals (r), where values greater than 0.5 are deemed highly-related pairs (i.e. parent-offspring), and values lower than 0.25 indicate unlikely-related pairs.

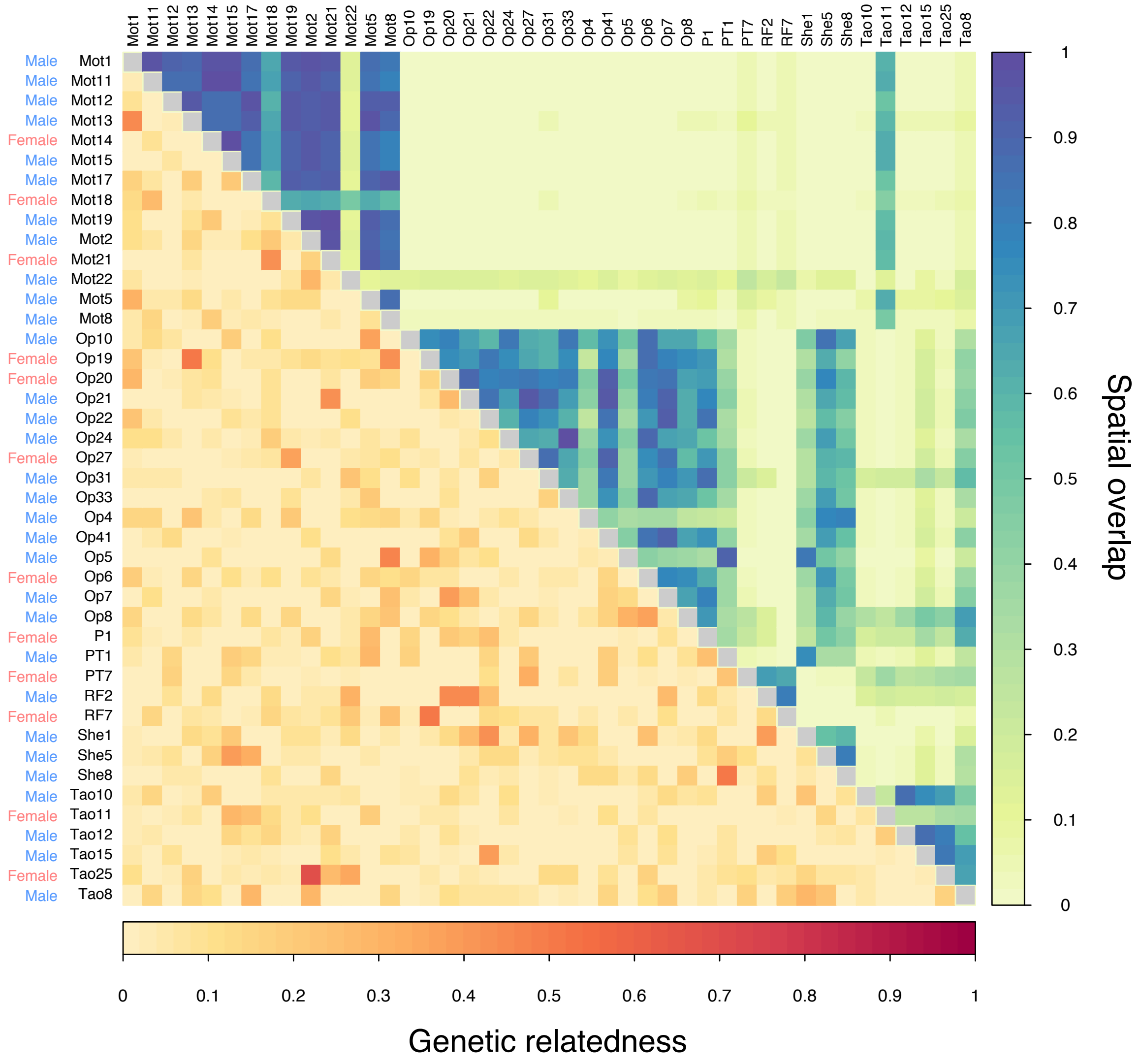

**References:**

Godde S, Humbert L, Côté SD, Réale D, Whitehead H. 2013. Correcting for the impact of gregariousness in social network analyses. Anim Behav. 85(3):553–558. doi:10.1016/j.anbehav.2012.12.010.

Milligan BG. 2003. Maximum-Likelihood Estimation of Relatedness. Genetics. 163(3):1153–1167.

Mourier J, Planes S. 2013. Direct genetic evidence for reproductive philopatry and associated fine-scale migrations in female blacktip reef sharks (Carcharhinus melanopterus) in French Polynesia. Mol Ecol. 22(1):201–214. doi:10.1111/mec.12103.

Queller DC, Goodnight KF. 1989. Estimating Relatedness Using Genetic Markers. Evolution. 43(2):258–275. doi:10.1111/j.1558-5646.1989.tb04226.x.

Wang J. 2002. An Estimator for Pairwise Relatedness Using Molecular Markers. Genetics. 160(3):1203–1215.

Wang J. 2007. Triadic IBD coefficients and applications to estimating pairwise relatedness. Genet Res. 89(3):135–153. doi:10.1017/S0016672307008798.
